# Dihydro-alpha-lipoic acid binds to human serum albumin at Sudlow I binding site

**DOI:** 10.1101/2020.10.16.342121

**Authors:** Nikola Gligorijević, Vladimir Šukalović, Goran Miljuš, Olgica Nedić, Ana Penezić

**Author notes:** Corresponding author: Ana Penezić, Institute for the Application of Nuclear Energy, Department for Metabolism, University of Belgrade, Banatska 31b, 11080 Belgrade.

## Abstract

Binding of dihydro-alpha-lipoic acid (DHLA) to human serum albumin (HSA) was characterised in detail in this study. Binding process was monitored by spectroscopic methods and molecular docking approach. HSA binds DHLA with moderate affinity, 0.80 ± 0.007 × 10^4^ M^−1^. Spectroscopic data demonstrated that the preferential binding site for DHLA on HSA is IIA (Sudlow I). Hydrogen bonds and electrostatic interactions were identified as the key binding interactions. DHLA binding thermally stabilized HSA, yet it had no effect on HSA structure and its susceptibility to trypsin digestion. Molecular docking confirmed that Sudlow I site accommodated DHLA in a certain conformation in order for binding to occur. Molecular dynamic simulation showed that formed complex is stable. Reported results offer future perspectives for investigations regarding the use of DHLA as a dietary intervention but also raise concerns about the effectiveness of alpha-lipoic acid and DHLA in treatment of patients with COVID-19.

## INTRODUCTION

Human serum albumin (HSA) is the most dominant protein in the circulation, with a referent concentration range from 35 to 50 g/L (522 μM to 746 μM). This is a protein with molecular mass of 67 kDa (Wang, Tian, & Chang, 2012). Structurally, HSA is composed of three homologous domains (I, II and III), each divided into two subdomains, A and B. Dominant secondary structure motif of HSA is α-helix (McLachlan & Walker, 1977).

HSA has many important functions in the circulation. Due to its high concentration, HSA participates in the osmotic pressure regulation (Lee & Wu, 2015). Its free Cys34 thiol group (in healthy individuals 70–80% of Cys34 thiol group is in a reduced form), makes HSA an important factor for plasma antioxidant capacity, contributing by 80% to the total plasma thiol amount (Pavićević et al., 2014). HSA is also a general transporter of fatty acids, ions and drugs. Due to its structure, HSA is able to accommodate and bind a variety of small molecules with moderate to high affinities. Two main binding sites for a plethora of different molecules (excluding fatty acids) are located at IIA subdomain or Sudlow I binding site, and IIIA subdomain or Sudlow II binding site. Drugs warfarin and ibuprofen are stereotypical ligands for Sudlow site I and Sudlow site II, respectively (Fasano et al., 2005).

Lipoic acid (LA) is a naturally occurring molecule whose main sources are potato, broccoli and spinach. Humans can also synthetize LA in small amounts. LA is readily absorbed from foods and its oral administration as a drug is a viable therapeutic option, including the treatment of patients with COVID-19 infection (Horowitz & Freeman, 2020; Zhang & Liu, 2020). LA supplements are also commercially available, with LA concentrations up to 600 mg per tablet. LA is shown to improve glycemic control, alleviate symptoms of diabetic polyneuropathy and is also effective against toxicity caused by heavy metal poisoning. Antioxidant activity of LA is manifested through ROS scavenging, transition metal ions (e.g., iron and copper) chelating, cytosolic glutathione and vitamin C levels increase, and oxidative stress damage repair (Zuliani & Baroni, 2015).

Following cellular uptake, LA is reduced to dihydrolipoic acid (DHLA), which is a very potent reducing agent (Zuliani & Baroni, 2015). LA has several beneficial effects such as antioxidant, improvement of glycemic control, mitigation of toxicity by heavy metal poisoning and immunomodulatory effects (Salinthone, Yadav, Bourdette, & Carr, 2008; Smith, Shenvi, Widlansky, Suh, & Hagen, 2004; Zuliani & Baroni, 2015).

Although the ability of albumin to bind DHLA is well known (Kawabata & Packer, 1994), no detailed analysis of this interaction has been reported so far. In the case of bovine albumin (BSA), DHLA was shown to bind at IIIA site (Suji et al., 2008), however no binding experiments in the presence of the specific ligand for this site were performed. Taking into account structural similarity of DHLA and octanoic fatty acid, it was proposed that DHLA binds to IIA site (Atukeren, Aydin, Uslu, Gumustas, & Cakatay, 2010), however, IIIA site was also considered (Suji et al., 2008).

Having in mind that DHLA is a very potent antioxidant and its use can alleviate a number of conditions related to oxidative stress, it seemed relevant to elucidate its mode of interaction with HSA, a universal transporter in the circulation. The properties of this interaction, are still unknown and undefined, so the present study aimed to investigate characteristics of the DHLA-HSA binding in detail, by using spectroscopic and molecular docking approach.

## MATERIALS AND METHODS

### Materials

All chemicals used were of analytical grade and were purchased from Sigma (Burlington, Massachusetts, USA). Stock solution of HSA, purchased from Sigma (A-1653) and used without additional purification, was made by dissolving HSA in 10 mM PBS, pH 7.4. The concentration of HSA was determined by using bicinchoninic acid (BCA) assay kit (Thermo Fisher Scientific, Waltham, Massachusetts, USA). Stock solution (5 mM) of DHLA was prepared by suspending DHLA in 10 mM PBS and then adding a small volume of 1 M NaOH until full clarification of solution was reached (Perricone et al., 1999). Trypsin was purchased from the Institute Torlak (Belgrade, Serbia) as a 0.25 % solution. All experiments were performed in triplicate at room temperature, using 10 mM PBS, pH 7.4, unless otherwise stated.

### Spectrofluorometric analysis of HSA-DHLA complex formation

Binding constant (Ka) of HSA-DHLA complex was determined by recording the quenching of intrinsic fluorescence emission of HSA (0.4 μM) in the presence of increasing concentrations of DHLA (from 4 to 35 μM) at 37 °C. Fluorescence spectra were recorded using FluoroMax®-4 spectrofluorometer (Horiba Scientific, Japan). HSA was exciteed at 280 nm and emission spectra were recorded in the range from 290 to 450 nm. Each spectrum was corrected for the emission of the control that contained only DHLA at particular concentration. The change of the emission intensity at 338 nm (HSA emission maximum) was used for the calculation of the binding constant. Emission intensity measured for HSA was first corrected for the small inner filter effect of DHLA using the equation:

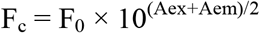

where Fc is corrected fluorescence, F_0_ is measured fluorescence, Aex and Aem are absorbances at excitation and emission wavelengths which are 290 nm and 338 nm, respectively.

Using corrected fluorescence, binding constant between HSA and DHLA was calculated using the following equation:

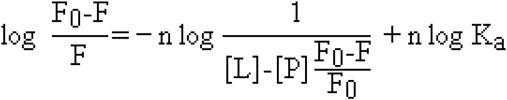

where F_0_ and F represent intensities of emission signals of HSA in the absence and in the presence of DHLA, [L] represents the total concentration of ligand (DHLA) and [P] the total concentration of protein (HSA).

Type of quenching, whether it’s static (complex formation) or dinamic, was determined by ploting Stern-Volmer (SV) graph and calculating SV quenching constant (Ksv) from it by applying the following equation:

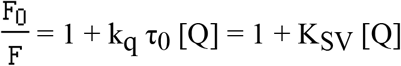

where F_0_ and F are intensities of emission signals without and in the presence of DHLA, k_q_ represents the biomolecule quenching rate constant, τ0 is the average lifetime of the biomolecule without quencher (10^−8^ s), [Q] is the total concentration of quencher (DHLA). The slope from SV plot represents K_SV_. K_SV_ was further used for the calculation of k_q_.

Thermodynamic parameters of DHLA binding to HSA were calculated by using the same experimental approach as for Ka calculation but at three different temperatures, 25, 30 and 37 °C. Calculated binding constants at three temperatures were then used to plot Van’t Hoff graph. Enthalpy (ΔH) and entropy (ΔS) were calculated from that graph applying the following equation:

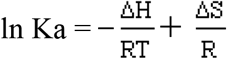

where T is temperature in Kelvins (K) and R is a universal gas constant (8.314 Jmol^−1^K^−1^). ΔH was calculated from the slope of Van’t Hoff graph and ΔS from the intercept. The change in Gibbs free energy was calculated from the equation:

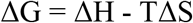

For specific fluorescence emission changes of 18 Tyr residues or the only Trp214 residue, synchronous fluorescence spectra were recorded on RF-6000 spectrophotometer (Shimadzu, Japan). Spectra were recorded in the range from 280 to 330 nm with Δλ of 60 nm for Trp214 and in the range from 290 to 325 nm with Δλ of 15 nm for Tyr residues. Here, Δλ represents Δλ of emission – Δλ of excitation for each specific residue.

For the confirmation of the specific binding site for DHLA on HSA, site IIA (Sudlow I) on HSA (0.4 μM) was blocked using site-specific ligand warfarin (40 μM). DHLA (20 and 40 μM) was added to this mixture and specific fluorescence emmision of wafarin (λex = 310 nm) was recoreded in the range from 340 to 440 nm (Vasquez, Vu, Schultz, & Vullev, 2009).

### Circular dichroism (CD) spectropolarimetric analysis of HSA-DHLA complex

The influence of DHLA binding on HSA structure was determined by CD-spectropolarimeter J-815 (Jasco, Japan) at room temperature and scan speed of 50 nm/min. Different concentrations of DHLA were added (6, 15 and 30 μM) to HSA (3 μM). Both HSA and DHLA stock solutions were dissolved in 10 mM phosphate buffer, pH 7.4. Tertiary protein structure was analyzed by recording near-UV CD spectra in the range from 260 to 320 nm using a cell path of 10 mm, while secondary protein structure was monitored by recording a far-UV CD spectra in the range from 185-260 nm using a cell path of 0.5 mm. Spectra obtained for mixtures were corrected for spectra derived from DHLA alone.

### UV-VIS analysis of HSA-DHLA complex

UV-VIS spectra of HSA (9 μM) in the presence of DHLA at different concentrations (9, 45 and 90 μM) were recorded at room temperature using Ultrospec 2000 spectrophotometer (Pharmacia Biotech, Sweden) in the range from 250 to 300 nm. A spectrum of each mixture was corrected for a spectrum obtained for DHLA alone. Also, UV-VIS spectrum of DHLA (90 μM) in the presence of HSA (9 μM) was recorded in the range from 300 to 450 nm and corrected for a spectrum obtained for HSA alone.

### Temperature stability analysis of HSA-DHLA complex

Temperature stability of HSA (0.4 μM) alone and in the presence of DHLA (40 μM) was determined by recording the reduction of fluorescence emission at 338 nm (emission peak of HSA) and at 335 nm (emission peak of HSA-DHLA complex), using the same equipment as in the titration experiment. Emission was recorded in the temperature range from 37 to 87 °C with a temperature increase rate of 2 °C. A mixture was allowed to equilibrate for 1 min before the measurement at each temperature. The obtained spectra were corrected by subtracting spectra of DHLA alone at each temperature. Results were fitted to sigmoid curves where inflection points represent melting temperatures (Tm).

### Proteolytic analysis using trypsin

For the investigation if DHLA binding affects susceptibility of HSA to trypsin proteolysis, the following experiment was performed at 37 °C. To solutions containing 4 μM HSA, alone and in the presence of DHLA (40 μM), 25 μL of 0.25 % trypsin solution was added. The final volume of reaction mixtures was 1 mL. At certain time points (1, 5, 10, 20 and 30 min) 50 μL aliquots were taken from the reaction mixture and PMSF immediately added at the final concentration of 2 mM, thus stopping the reaction. Proteolytic fragments of HSA were analyzed by reducing SDS-PAGE using a 12 % gel in a standard manner. Gel was stained using Silver Stain Plus Kit (Bio-Rad, Hercules, California, USA).

### Docking simulations

Docking simulations were carried out with Schrodinger Maestro Suite (Schrödinger, LLC, New York, NY, 2018) using crystal structure of HSA complexed with warfarin (PDB code: 2BXD, (Ghuman et al., 2005), obtained from RCSB PDB database (https://www.rcsb.org/). DHLA structure was drawn in ChemDraw program (PerkinElmer Informatics, 2017). All structures were prepared in Maestro software, using default procedures. Up to 20 different docked structures were generated with Induced fit docking protocol (Sherman, Day, Jacobson, Friesner, & Farid, 2006). The obtained docking structures were examined and the best structure was selected based on the number of receptor-ligand interactions and docking score.

### Molecular dynamics (MD) simulations

MD simulations were done in Schrodinger Desmond software package (Bowers et al., 2006). Selected docked structure for MD was solvated with TIP3P explicit water model, and neutralized via counter ions. Salt solution of 0.15 M KCl was added. To calculate the interactions between all atoms OPLS 2003 force field was used. For the calculation of the long-range Coulombic interactions, particle-mesh Ewald (PME) method was used, with the cut-off radius of 9 Å for the short-range Van der Waals (VdW) and electrostatic interactions.

During the course of the simulation, constant temperature of 310 K and a pressure of 1.01235 bar were maintained, using the Nose–Hoover thermostat, and the Martyna Tobias Klein method. 50 ns MD simulation with 2.0 fs step was performed and the collected trajectory used in the MD analysis to asses docking pose and the stability of protein-ligand interactions.

## RESULTS AND DISSCUSION

### Binding of DHLA by HSA

The presence of DHLA quenches intrinsic fluorescence of HSA, as can be seen from Figure 1A. Moreover, a very small blue shift of 3 nm is observed at the emission maximum of HSA, as the concentration of DHLA increases. These results suggest that HSA binds DHLA and that polarity of the surrounding of Tyr and Trp214 amino acid residues is not significantly altered. Fluorescence quenching can be both dynamic and static (complex formation). In order to determine which type is present here, SV graph was plotted (Figure 1B) and from its slope Ksv was calculated. The obtained SV plot is linear (r^2^ = 0.99), indicating that only one type of quenching occurs in the observed system. Ksv was calculated to be 0.83 × 10^4^ M^−1^ and the quenching rate constant of the biomolecule, k_q_, was calculated to be 0.83 × 10^12^ M^−1^. Since k_q_ is about two orders of magnitude higher than the diffusion rate of biomolecules (10^10^ M^−1^s^−1^), this result strongly suggests the presence of static type of quenching, confirming that HSA binds DHLA. From the equation (2) and the plot from Figure 1C, Ka at 37 °C was calculated to be 0.80 ± 0.007 × 10^4^ M^−1^, showing that HSA binds DHLA with moderate affinity.

**Figure 1.**
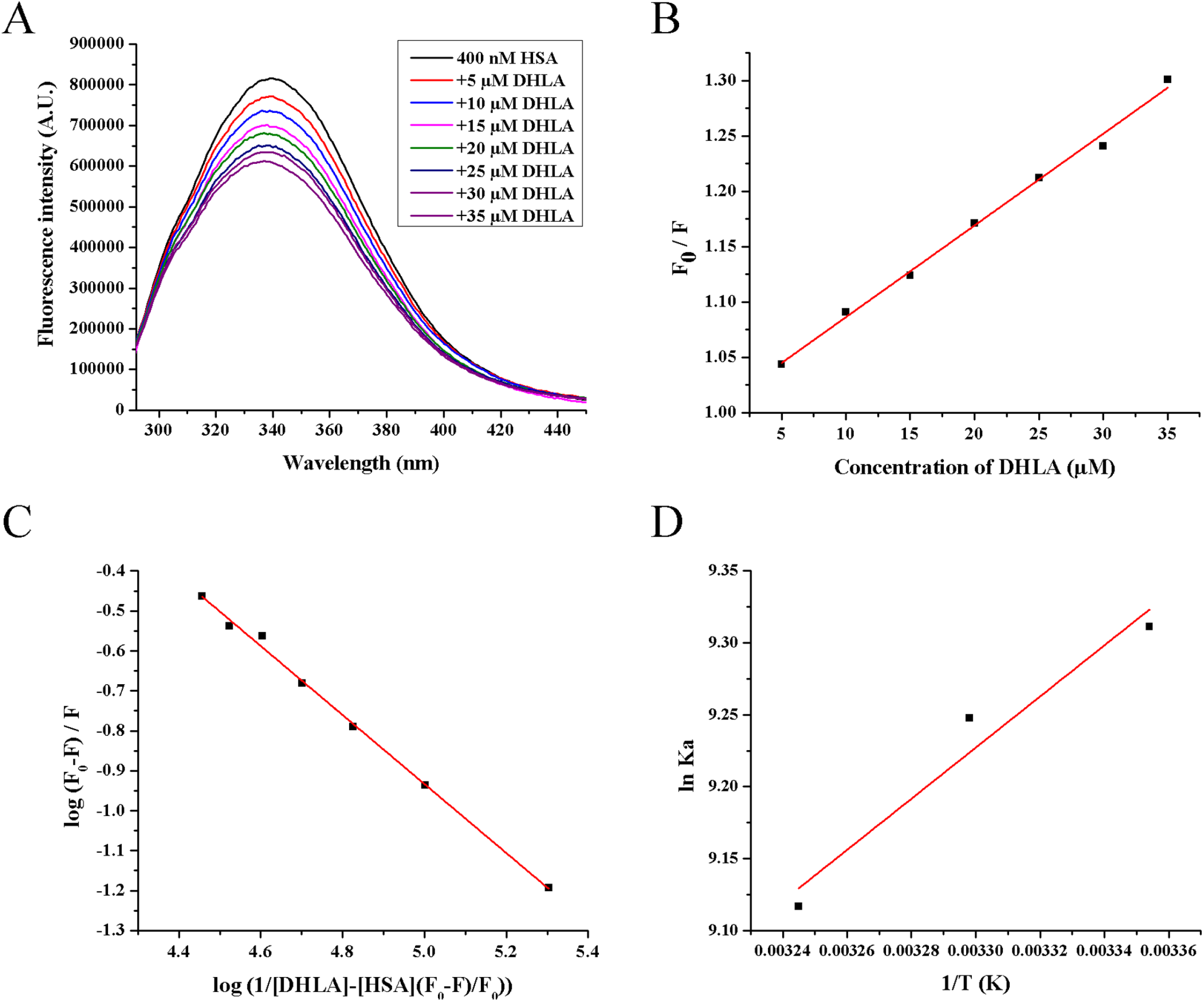
Binding analysis of HSA and DHLA using spectrofluorimetry. Fluorescence emission spectra of HSA (excited at 280 nm) in the presence of increasing concentrations of DHLA (A). Stern-Volmer plot (B) and plot used for the determination of the binding constant between HSA and DHLA (C) obtained using fluorescence emission maximum of HSA at 338 nm. Van’t Hoffs graph obtained by calculating the binding constant between HSA and DHLA at three different temperatures (D).

When Ka was calculated at three different temperatures, its value decreased as the temperature increased. This usually occurs, but is not exclusive, in static type of fluorescence quenching (excluding entropy driven binding) since complex formation is weaker at higher temperatures (Van De Weert & Stella, 2011). Using the obtained Ka values at three temperatures, thermodynamic parameters were calculated from Van’t Hoff plot (Figure 1D). Large negative value of ΔH was obtained, −32 kJmol^−1^ as well as small negative value of ΔS, 29 Jmol^−1^K^−1^. These results indicate that electrostatic interactions, hydrogen bonds and Van der Walls interactions are mainly responsible for complex formation between HSA and DHLA. The change in Gibbs free energy, ΔG, at 37 °C was calculated to be −23 kJmol^−1^.

Synchronous fluorescence spectra can give information about changes in the emission of Tyr and Trp amino acid residues. Since HSA has only one Trp residue, located inside the binding site IIA (Sudlow I) (Salem, Lotfy, Amin, & Ghattas, 2019), information from synchronous spectra provides insight into the binding place for certain ligand. In the presence of increasing concentrations of DHLA, Trp specific emission spectrum was significantly quenched (Figure 2A), while that originating from Tyr was reduced to a very small extent (Figure 2B). Considering the position of the Trp residue in HSA, this result strongly indicated that the binding site for DHLA is located in IIA subdomain (Sudlow I). To confirm this, DHLA was added to HSA in the presence of warfarin, and the change in warfarin fluorescence was recorded. When bound to HSA, warfarin fluorescence intensity increases at its emission maximum (Figure 2C). This is a usual consequence of ligand binding to a protein, since the ligand becomes shielded from water and located in a more hydrophobic environment (Liang, Tajmir-Riahi, & Subirade, 2008). Similar observation was recorded in the case of phycocyanobilin (PCB) binding to HSA that occurs at both IIA and IB subdomains of HSA (Minic et al., 2015). Binding of warfarin to HSA is well characterized with the affinity constant of about 10^5^ M^−1^ (Li et al., 2014). As warfarin specifically binds to Sudlow I site on HSA, it is used to block this site in the studies aimed to locate the exact binding site for other ligands (Petitpas, Bhattacharya, Twine, East, & Curry, 2001). Results shown in Figure 2C indicate that the emission spectrum of warfarin remains the same in the presence of DHLA (at two concentrations), confirming that binding site IIA on HSA remains occupied by warfarin, thus suggesting that this site is the preferential binding site for DHLA. Even in equimolar concentrations of DHLA and warfarin, fluorescence spectrum of warfarin remains unaltered, indicating that HSA binding affinity for DHLA is lower than for warfarin, which is in agreement with the calculated Ka for HSA-DHLA complex. Having this in mind, it is noteworthy to mention a current pandemic situation with COVID-19 and its potential treatment with alpha lipoic acid (Horowitz & Freeman, 2020; Zhang & Liu, 2020). It was proposed that LA blocks NF-κB and cytokine formation, and thus alleviates cytokine storm syndrome in critically ill patients (Horowitz & Freeman, 2020). Since warfarin and its derivatives are commonly used as anticoagulants (also included in the therapy of severe cases with COVID-19) which preferentially bind to Sudlow I site on HSA, it is questionable whether LA treatment of patients infected with virus Sars-CoV-2 is sufficiently beneficial if they also receive warfarin.

**Figure 2.**
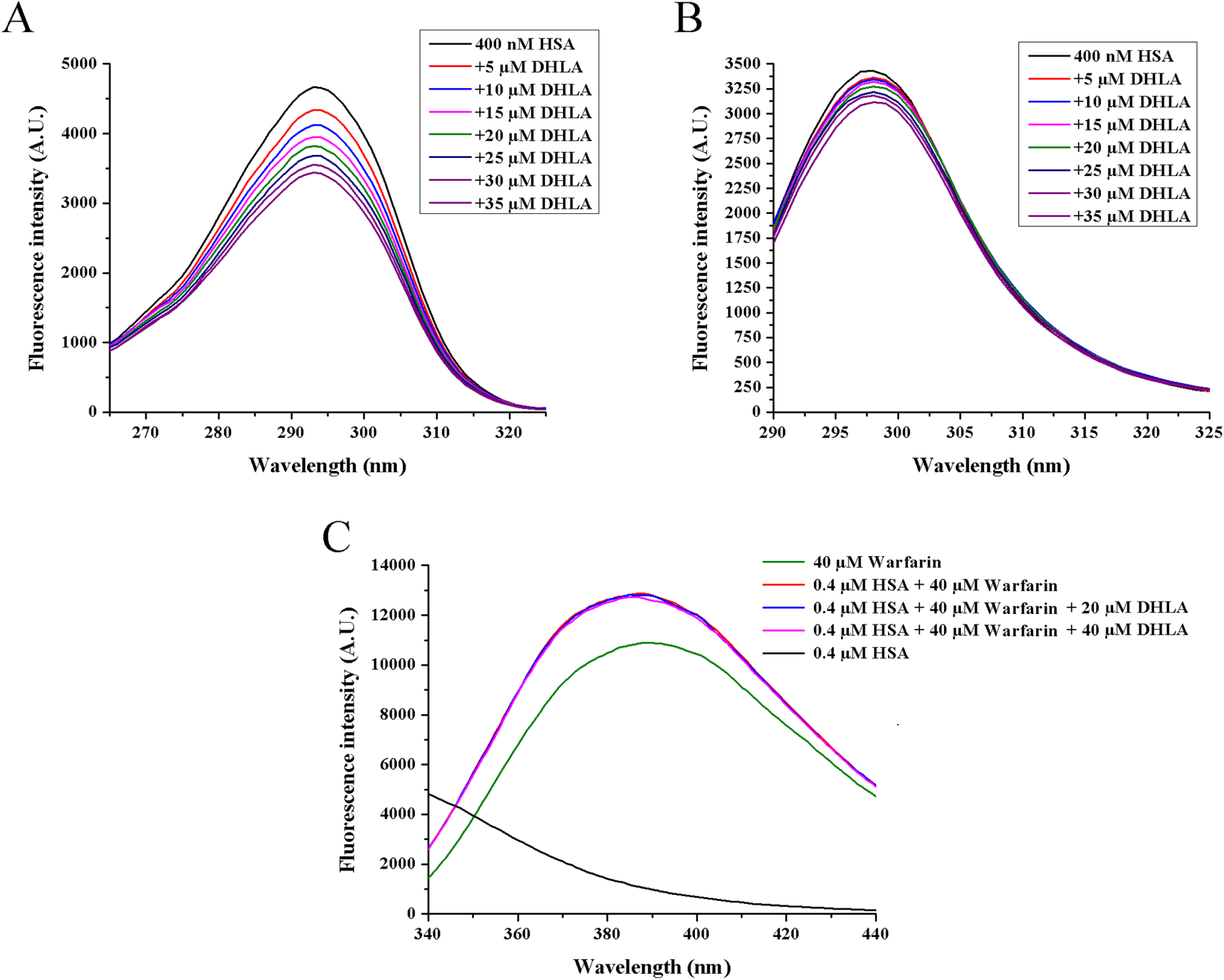
Determination of a binding site of DHLA on HSA. Synchronous fluorescence spectra of HSA with Δλ = 60 nm for Trp (A) and Δλ = 15 nm for Tyr (B) in the presence of increasing concentrations of DHLA. Fluorescence emission spectra of warfarin (excited at 280 nm) in the absence and in the presence of HSA, as well as in the presence of HSA and DHLA at warfarin/DHLA molar ratios of 2/1 and 1/1 (C).

Proteins absorb light in the UV region at about 280 nm due to the presence of aromatic amino acid residues and changes in the absorption spectrum of a protein in this region may occur as a consequence of changes in the polarity of the environment close to these residues. Figure 3A shows that the UV-VIS absorption spectrum of HSA does not change in the presence of increasing concentrations of DHLA, indicating that no significant conformational change of HSA occurs, thus confirming the results obtained by spectrofluorimetry (Figure 3A). On the other hand, the absorption spectrum of DHLA shows both blue shift of its peak and the reduction of its intensity in the presence of HSA (Figure 3B). Similar effect was previously observed upon DHLA binding to fibrinogen (Gligorijević, Šukalović, Penezić, & Nedić, 2020) and upon binding of 2–amino-6-hydroxy–4–(4-N,N-dimethylaminophenyl)-pyrimidine-5-carbonitrile to BSA (Suryawanshi, Walekar, Gore, Anbhule, & Kolekar, 2016). Change in the absorption spectrum of DHLA is an additional proof that it forms a complex with HSA.

**Figure 3.**
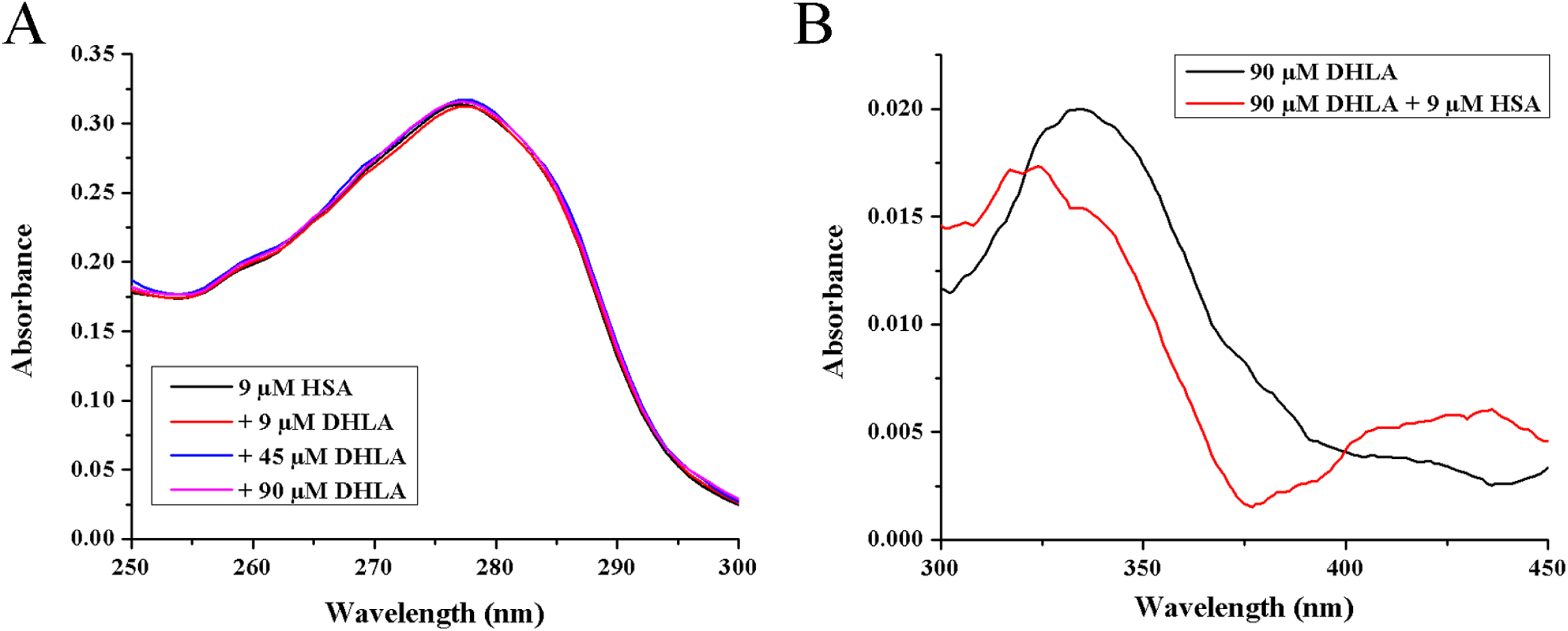
Analysis of structural alterations of HSA and DHLA due to mutual binding. Far-UV CD (A) and near-UV CD (B) spectra of HSA alone and in the presence of increasing concentrations of DHLA. UV absorption spectra of HSA alone and in the presence of increasing concentrations of DHLA (C). UV-VIS absorption spectra of DHLA alone and in the presence of HSA (D).

Protein structure is often affected by ligand binding. Some ligands induce more ordered structure, others more disordered, and some have no effect. HSA contains only α-helixes as elements of the secondary structure. It was shown that binding of PCB and amoxicillin to HSA increase its content of α-helixes (Radibratovic et al., 2016; Yasmeen, Riyazuddeen, & Rabbani, 2017), while binding of plumbagin, safranal and crocin decrease it (Qais, Husain, Khan, Ahmad, & Hassan, 2020; Salem et al., 2019). The obtained far-UV CD spectra of HSA (Figure 4A) show a typical signal of the protein where α-helixes are dominant, with characteristic negative wide peak in the range from 209 to 220 nm. As it can be seen from this figure, no significant change in the secondary structure of HSA occurs upon binding of DHLA, even when the concentration of DHLA is ten times larger than HSA. Tertiary structure of HSA is also unaltered due to DHLA binding since near-UV CD spectra are practically the same in pre presence of all tested DHLA concentrations (Figure 4B).

**Figure 4.**
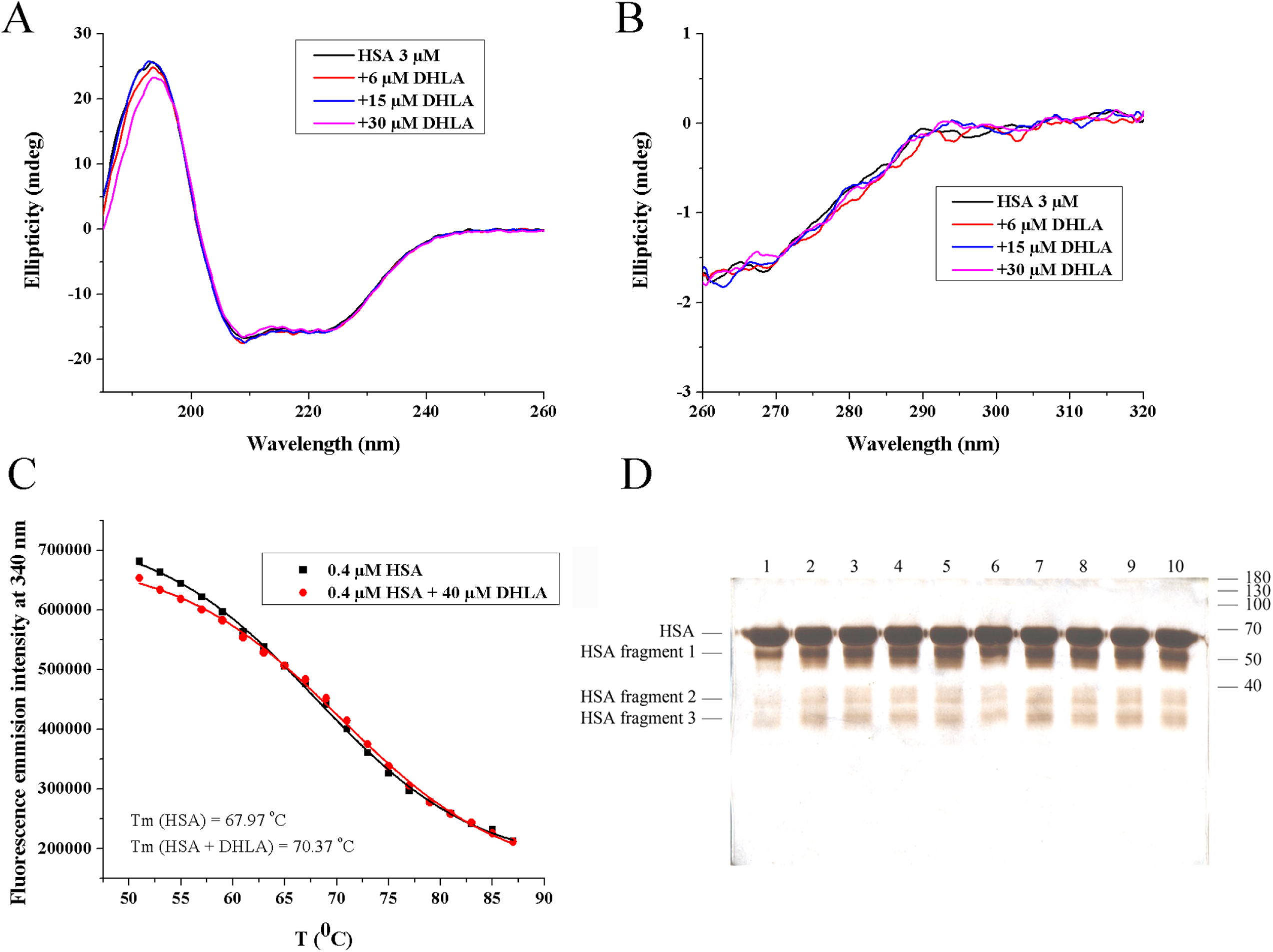
Analysis of temperature stability of HSA alone and in the presence of DHLA (A). Analysis of HSA digestion by trypsin in the absence (lanes 1-5, samples taken after 1, 5, 10, 20 and 30 min of proteolysis) and in the presence of DHLA (lanes 6-10) by reducing SDS-PAGE on 12 % gel (B).

### Stability of HSA-DHLA complex

Factors that may affect melting point of a protein, besides its structure, include the presence of bound molecules as well as their structure. When a complex forms, new interactions establish that may contribute to altered thermal stability of a protein. In the case of HSA, free protein has Tm of approximately 62 °C, while bound fatty acids increase its thermal stability reaching Tm from 64 to 72 °C (Lang & Cole, 2015). Certain ligands, such as PCB and embelin, also increase thermal stability of HSA (Radibratovic et al., 2016; Yeggoni, Rachamallu, & Subramanyam, 2016). On the contrary, some drugs, such as amoxicillin, decrease thermal stability of HSA upon binding (Yasmeen et al., 2017). Commercial HSA used in this study had Tm of 68 °C. In the presence of DHLA, Tm of HSA increases to 70 °C (Figure 4C). Even though DHLA didn’t change the structure of HSA significantly upon binding (Figures 4A and 4B), it seems that new interactions in this complex additionally thermally stabilized the protein.

Increased Tm of HSA in the presence of DHLA indicates that rigidness in the protein structure increases, which may affect its susceptibility to proteolytic cleavage. In order to be proteolyzed, peptide bonds in the protein need to be flexible and exposed enough to enable its accurate accommodation in the active site of a protease. Some ligands, such as bilirubin, reduce the susceptibility of HSA to cleavage by trypsin (Sjödin, Hansson, & Sjöholm, 1977). According to the results of this study, it seems that DHLA, although it thermally stabilizes HSA, has no significant effect on HSA proteolysis by trypsin (Figure 4D). Thus, it may be expected that the formation of HSA-DHLA complex will not have significant (if any) effect on the protein half-life in circulation in respect to proteolysis. The first and the dominant fragment of HSA resulting from proteolysis by trypsin is the one at about 50 kDa, while other fragments with lower concentrations and molecular masses appear later. This finding is in accordance with the already published data (Radibratovic et al., 2016).

### Molecular modeling

Binding site IIA consists of a binding pocket deeply embedded in the core of the subdomain that is formed by all six helices of the subdomain and the loop-helix residues 148-154 of IB (Ghuman et al., 2005). Pocket interior is predominantly hydrophobic, apart from the two clusters of polar residues (Tyr150, His242, Arg257 and Lys195; Lys199, Arg218 and Arg222).

Induced fit docking simulation results have shown that DHLA binds to HSA BS II site (Figure 5). The energetically most favorable conformation of the docked pose has showed that the key interactions are salt bridges formed by DHLA carboxyl group with Arg18 and Arg222 of HSA, followed by hydrogen bonds formed between DHLA sulfhydryl group and Arg257, Ser287 (Figure 5). Molecular docking analysis suggested that DHLA binds at Sudlow I site in a defined conformation, thus favoring interactions with specific amino residues. Having in mind that DHLA has high torsional flexibility due to nine dihedral angles which give many possible rotamers (Vigorito, Calabrese, Paltanin, Melandri, & Maris, 2017), a recorded change in the absorption spectrum (Figure 3B) could point to a DHLA-conformational shift towards rotamers with the highest probability of being bound to HSA.

**Figure 5.**
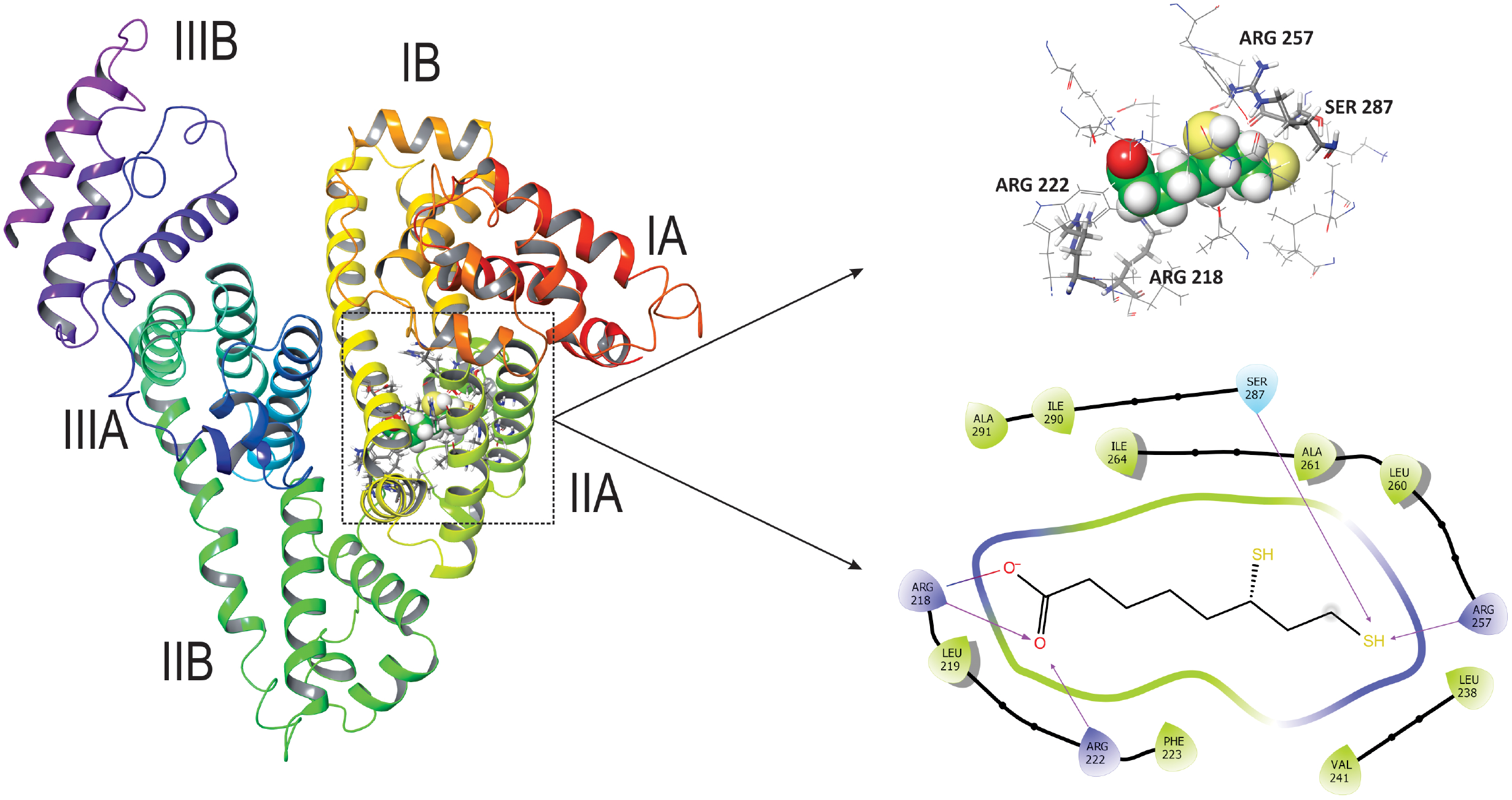
An overview of HSA with DHLA docked into BS II. Domains are color coded and represented as secondary structure ribbons. BS II composition and key interactions diagram. All amino acid residues in close contact with DHLA are displayed, with key amino acid residues marked.

To verify docking simulation results, DHLA-HSA interactions were monitored throughout 50 ns molecular dynamic simulation. MD starting point was the best conformation obtained in docking stage. The obtained MD trajectory was analyzed both in terms of complex stability and the persistence of key DHLA-HSA interactions over simulation time period.

The observed RMSD values for HSA show that simulation has equilibrated, fluctuations fall within 1 – 2.5 Å range suggesting minor conformational changes during simulation (Figure 6A).

**Figure 6.**
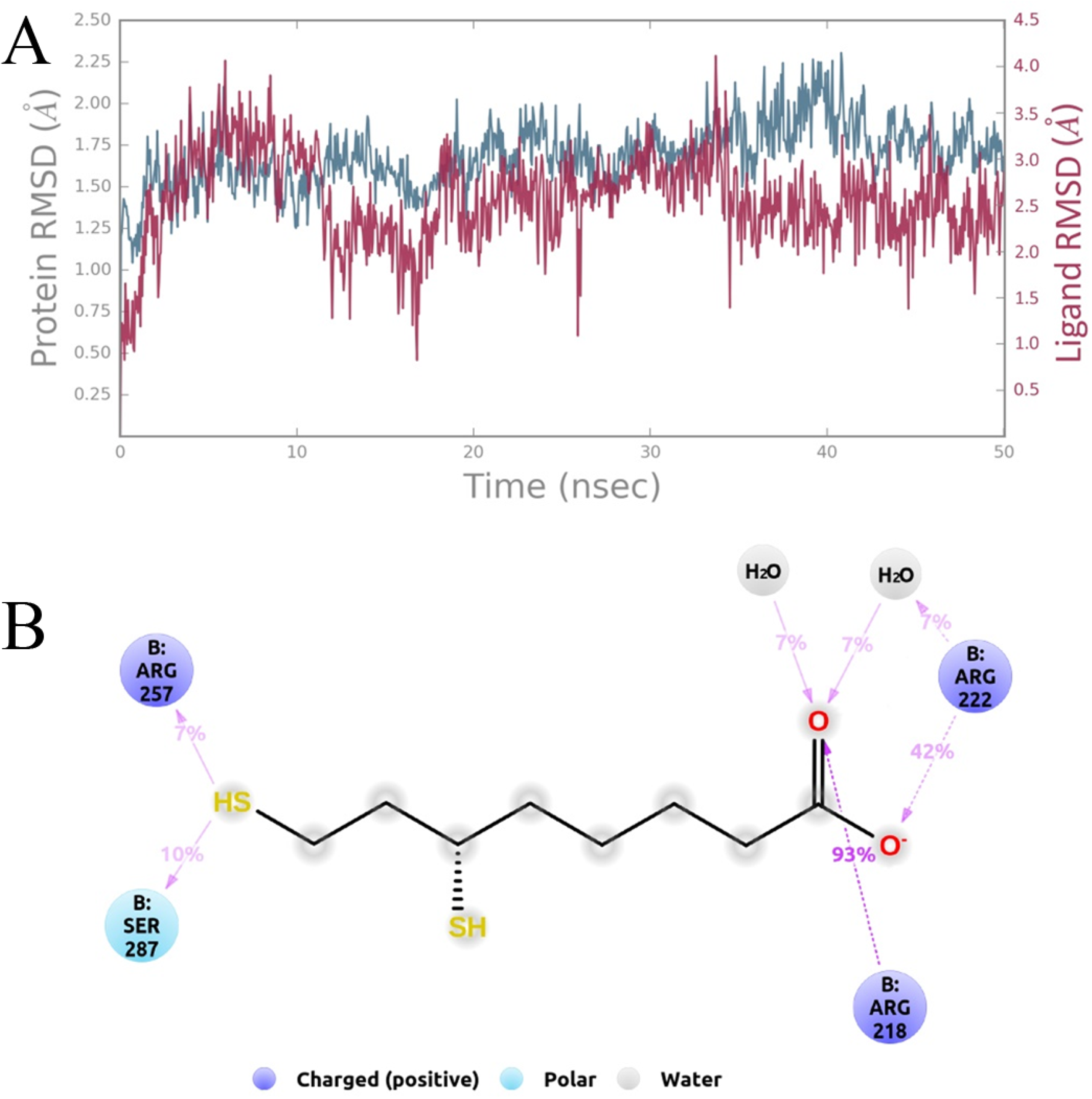
HSA and DHLA RMSD plot (A) and the observed key interactions during 50 ns simulation time (B).

Monitored HSA-DHLA interactions showed that key interaction is a salt bridge formed between DHLA carboxyl group and Arg218. This interaction is present over 93% of simulation time, making it crucial for DHLA binding to HSA and it also indicates correct orientation of DHLA inside the BS. Salt bridge with Arg218 is reinforced by interaction with Arg222 (42% of simulation time). Once DHLA is in correct position in the BS, additional hydrogen bonds between sulfhydryl group and Ser287, Arg257 are established. Those hydrogen bonds are maintained for a 10% (Ser287) and 7% (Arg257) of total simulation time (Figure 6B). All other interactions are observed for less than 5% of total simulation time (Supplementary Figure 1).

HSA can be modified by oxidation, and slight structural alterations, resulting from this chemical modification, cause an impairment of HSA functions, including its ligand binding ability (Kawakami et al., 2006). Redox state of Cys34 on HSA can also influence its binding properties (Oettl & Stauber, 2007). Considering its high concentration and the capacity to bind wide range of drugs, changes in binding properties of HSA may have a significant impact on pharmacokinetic and pharmacodynamic (PKPD) characteristics of prescribed drugs. DHLA was shown to be able to protect serum albumin from glycation (Kawabata & Packer, 1994), methylglyoxal modification (Sadowska-Bartosz, Galiniak, & Bartosz, 2014) and it can protect Cys34 from oxidation (Atukeren et al., 2010). Thus, by binding to HSA, DHLA can directly protect HSA from oxidation and, at the same time, keep the binding and antioxidative properties of HSA unaltered. Since it is used as a food supplement, a detailed PKPD knowledge on DHLA is very important, including information on its binding partners in the circulation. Besides HSA, fibrinogen was also shown to bind DHLA with a similar affinity (Gligorijević et al., 2020).

## CONCLUSION

The obtained results describe in detail the binding of DHLA to HSA for the first time. Experimental results have shown that binding site IIA or Sudlow I is the preferential binding site for DHLA. Molecular docking analysis and dynamics confirmed the ability of Sudlow I site to accommodate DHLA and that the formed complex is stable. The binding of DHLA doesn’t alter significantly the structure of HSA, although it stabilizes the protein itself to some extent. HSA susceptibility to proteolytic cleavage by trypsin remains the same in the presence of DHLA, thus no change of HSA half-life in the circulation (regarding proteolysis) is expected. The reported results expand the knowledge on PKPD properties of DHLA and offer a future perspective for further investigations regarding the use of DHLA as a dietary intervention. Furthermore, the obtained results raise concerns whether alpha-lipoic acid and DHLA are sufficiently beneficial as a part of the proposed treatment protocol of patients with COVID-19 who are receiving warfarin therapy as well, due to their competitive binding and lower affinity of HSA for these antioxidants than for warfarin.

## Acknowledgments

This research was funded the Ministry of Education, Science and Technological Development of the Republic of Serbia, contract numbers: 451-03-68/2020-14/200019 and 451-03-68/2020-14/200026. There is no conflict of interest regarding this study.

---

**Supplementary Figure 1**. Summary of DHLA-HSA interactions observed during 50 ns simulation time. Each orange line represents one established interaction during 1 ns time frame.

